# Biological function polarity prediction of missense variants using machine learning

**DOI:** 10.1101/2020.04.03.023440

**Authors:** Adhideb Ghosh, Alexander A. Navarini

## Abstract

Functional interpretation is crucial when facing on average 20,000 missense variants per human exome, as the great majority are not associated with any underlying disease. *In silico* bioinformatics tools can predict the deleteriousness of variants or assess their functional impact by assigning scores, but they cannot predict whether the variant in question results in gain or loss of function at the protein level. Here, we show that machine learning can effectively predict this biological function polarity of missense variants. The new method adapts weighted gradient boosting machine approach on a set of damaging variants (1,288 loss of function and 218 gain of function variants) as annotated by the tools SIFT, PolyPhen2 and CADD. Area under the ROC curve of 0.85 illustrates high discriminative power of the classifier. Predictive performance of the classifier remains consistent against an independent set of damaging variants as highlighted by the area under the ROC curve of 0.83. This new approach may help to guide biological experiments on the clinical relevance of damaging genetic variants.

**Author summary:** Missense variant occurs when a single genetic alteration in DNA takes place and as a result a new amino acid is translated into the protein. This amino acid change can inactivate the existing protein function causing loss-of-function or produce a new function causing gain-of-function. Therefore, it is very important to interpret these functional consequences of missense variants as they often turn out to be disease causing. Each individual’s genome sequence has thousands of missense variants, out of which very few are actually associated with any underlying disease. Various computational tools have been developed to predict whether missense variants are damaging or not, but none of them can actually predict whether the damaging missense variants cause gain-of-function or loss-of-function. We have developed a new ensemble classifier to predict this biological function polarity at the protein level. The classifier combines the prediction scores of three existing bioinformatics tools and applies machine learning to make effective predictions. We have validated our classifier against an independent data set to show its high predictive power and robustness. The predictions made by our machine learning tool can be used as indicators of biological function polarity, but with further evidence on pathogenicity.

## Introduction

High-throughput sequencing technologies have allowed us to cost-efficiently screen patients’ genomes. The major goal behind most of this generated data is to discover disease-causing variants, which is the fundamental problem in medical genetics. Each individual genome has thousands of missense variants, but only few actually cause the underlying genetic diseases [1]. Hence, the main challenge lies in the understanding of functional consequences of these variants. Because of the sheer amount, it is not feasible for the researchers to experimentally examine all the missense variants for actual pathological consequences to investigate a potential disease-causing role.

Therefore, various *in silico* bioinformatics tools [2-9] have been introduced to predict the functional effect of genetic variants. Some tools perform their predictions based on the sequence conservation approach like SIFT [2], MutationAssessor [3], fathmm [4] and MutationTaster [5]. SIFT computes it from closely related sequences [2], MutationAssessor captures the evolutionary conservation patterns in a protein family and its subfamilies [3], fathmm combines the alignments with hidden Markov models [4] and MutationTaster uses Bayes classifier with three classification models to test different variant effects on the protein sequences [5]. PolyPhen2 [6] and CADD [7] integrate different information resources and combine them with a machine learning based approach. PolyPhen2 makes its prediction based on sequence, phylogenetic and structure based features [6]. CADD combines many diverse annotations into a single prediction score including allelic diversity, functionality, pathogenicity, disease severity, regulatory effects and associated complex traits [7]. As far as consensus approaches are concerned [8], Condel [9] calculates its score by combining the outputs of multiple tools including SIFT, PolyPhen2, MutationAssessor and fathmm. Although these tools can predict whether a missense variant is damaging or not, they cannot predict its actual biological consequences on the protein function. Flanagan et al. had tried to make such predictions using SIFT and PolyPhen prediction scores with a biased set of only 133 missense variants across three genes. But they ended up with very low specificity and poor accuracy in predicting variants that cause a functional change in the translated protein. [10]

The aim of our study was to identify a set of tools whose prediction scores can be utilized to predict the actual functional consequence of a missense variant, namely whether it causes gain of function (GOF) or loss of function (LOF) at the protein level. We have developed an ensemble classifier named BioPol (Biological function Polarity prediction of missense variants) for the same purpose by combining the predictions of different bioinformatics tools and taking variant class imbalance into consideration. BioPol was evaluated by appropriate performance metric and validated against an independent set of missense variants.

## Results

We assembled a total of 298 GOF variants and 1,583 LOF variants from ClinVar [11]. Only pathogenic and likely pathogenic variants were selected with at least one assertion criteria without any conflicts to ensure high confidence [11].

### Prediction score analysis of missense variants

The whole set of 1,881 high confidence variants was annotated using Ensembl web tool variant effect predictor (VEP) [12]. Out of more than 100 annotated columns, our primary focus was on the prediction score distribution of seven *in silico* bioinformatics tools (SIFT, PolyPhen2, Condel, CADD, fathmm, MutationAssessor and MutationTaster) across all pathogenic and likely pathogenic missense variants. We excluded the prediction scores of Condel, fathmm, MutationAssessor and MutationTaster from further analysis as they were unable to annotate at least 90% of GOF and LOF variants.

For the remaining three tools, we only considered the variants that were annotated by all of them. As a result a new set of 290 GOF and 1,537 LOF variants was obtained across 77 and 377 genes respectively (S1 Table). For each tool, we analyzed the prediction scores between GOF and LOF variants using a two-tailed unpaired Student’s t-test in order to check whether these scores could distinguish the variant classes or not. Furthermore, we refined this variant set to a new set of 218 GOF and 1,288 LOF variants, which were predicted to be damaging by all three tools to avoid any false positives. Prediction scores were statistically analyzed between two variant classes in a similar manner. Mean prediction scores for all tools are summarized in Table 1 for both refined variant data sets.

**Table 1.** Mean prediction score differences of SIFT, PolyPhen2 and CADD between GOF and LOF variants.

| *In silico*<br>bioinformatics<br>tools | Annotated ClinVar variants by<br>all tools |  |  | Predicted to be damaging ClinVar<br>variants by all tools |  |  |
| --- | --- | --- | --- | --- | --- | --- |
|  | GOF<br>(#<sup>a</sup> 290) | LOF<br>(#<sup>a</sup> 1,537) | P-value<sup>b</sup> | GOF<br>(#<sup>a</sup> 218) | LOF<br>(#<sup>a</sup> 1,288) | P-value<sup>b</sup> |
| Mean SIFT<br>score<sup>c</sup> | 0.052 | 0.023 | 0.005 | 0.007 | 0.003 | 1.59e<sup>-09</sup> |
| Mean PolyPhen2<br>score<sup>d</sup> | 0.81 | 0.87 | 0.0004 | 0.93 | 0.97 | 1.71e<sup>-06</sup> |
| Mean CADD<br>phred score<sup>e</sup> | 26.06 | 28.71 | $< 2.2e^{-16}$ | 27.46 | 29.73 | $< 2.2e^{-16}$ |

<sup>a</sup> # indicates number of variants in each class
<sup>b</sup> p values were calculated using two-tailed unpaired Student's t-test
<sup>c</sup> SIFT scores range from 0 to 1. Damaging variants: SIFT score $< 0.05$ .
<sup>d</sup> PolyPhen2 scores range from 0 to 1. Damaging variants: PolyPhen2 score $> 0.5$
<sup>e</sup> CADD phred scores range from 0 to 100. Damaging variants: CADD phred score $> 15$ .

We found the prediction scores of all tools to be significantly different (p < 0.01) between GOF and LOF classes as shown in Table 1. Mean prediction score differences between the classes become even more significant when we restrict the variants only to the predicted to be damaging ones. Therefore, prediction scores of all three tools were considered as potential biological function polarity predictors.

### BioPol algorithm

The combined set of 1,506 missense variants was randomly split into a training set (70%) and a test set (30%). We developed BioPol as a weighted gradient boosting machine [13] based classifier using R package caret [14]. SIFT, PolyPhen2 and CADD prediction scores were selected as the final set of predictors to assess whether a damaging missense variant is causing GOF or LOF at the protein level. We added equal weight of 0.5 to each variant class to resolve the class imbalance problem as the number of LOF variants is much higher than that of GOF variants by natural occurrence. For tuning model parameters, we repeated 10-fold cross validation 5 times and specified a tuning parameter grid. Average area under the ROC curve (AUROC) was used as the performance measure across 50 resamples to determine the optimal parameter set. The final classifier was built on the training set of 1,055 pathogenic missense variants using 450 trees with maximum tree depth of 3 and learning rate of 0.01.

### BioPol performance

ROC curve was plotted to measure the performance of BioPol, illustrating true positive rate as a function of false positive rate (Fig 1). An AUROC of 0.85 indicated high predictive power of the classifier on the test set of 451 pathogenic missense variants. This performance measure showed that BioPol can be used to predict GOF and LOF variants based on their SIFT, PolyPhen2 and CADD scores provided the tools predicted them as damaging.

**Fig 1.**
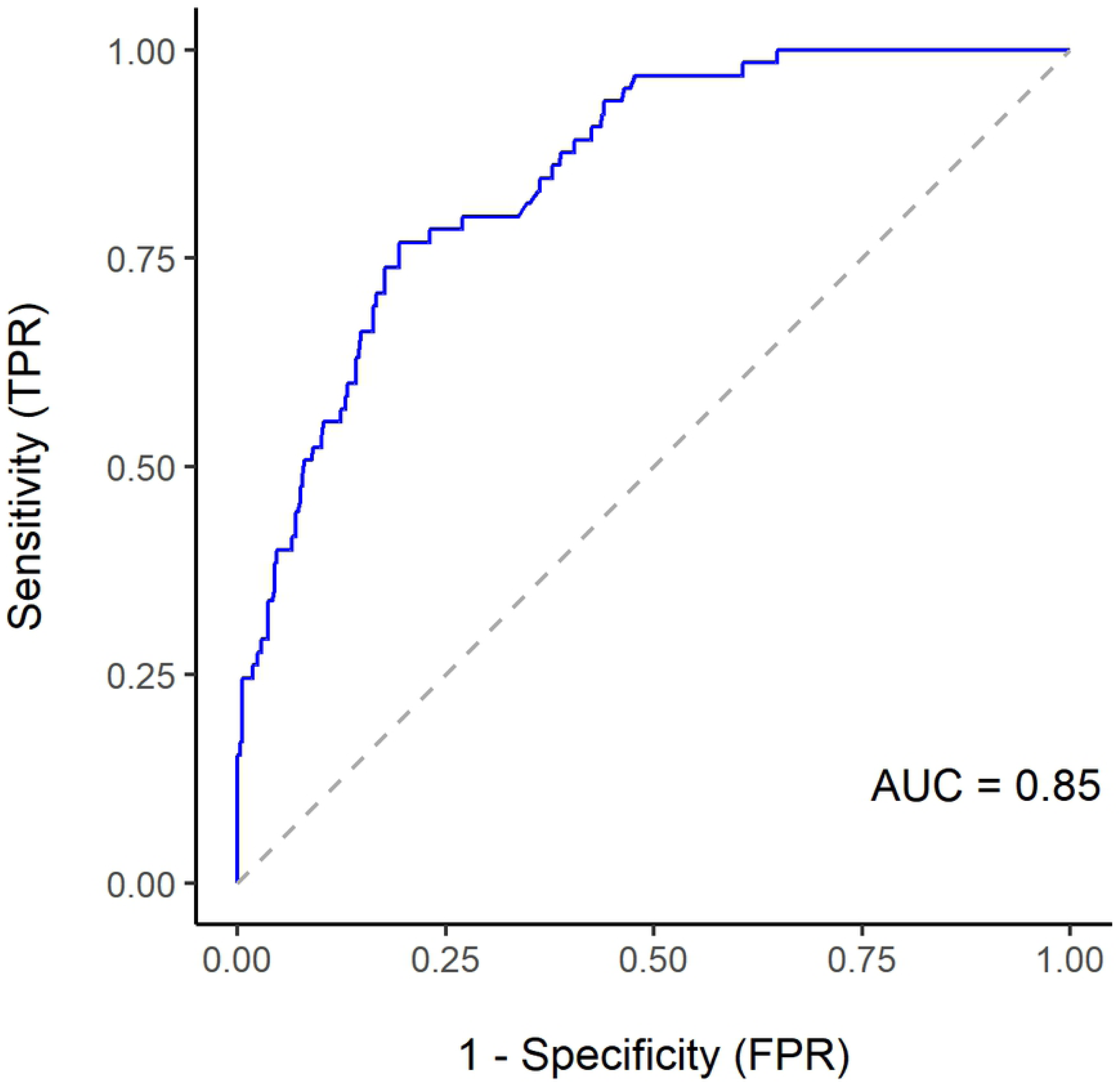
ROC curve illustrating predictive power of BioPol on the test set.

### BioPol validation

An independent high confidence validation set of pathogenic and likely pathogenic missense variants was constructed from ClinVar using a different submission time frame. We annotated them using VEP and restricted them to a set of 14 GOF and 80 LOF variants that were predicted to be damaging by all three tools. BioPol predictions on this validation set are summarized as a confusion matrix in Table 2 for both variant classes. ROC curve was made and an AUROC curve of 0.83 was found to highlight the predictive power of BioPol (Fig 2). BioPol predictions of 94 pathogenic missense variants from the validation set are available in the supplementary information for reference (S2 Table).

**Table 2.**
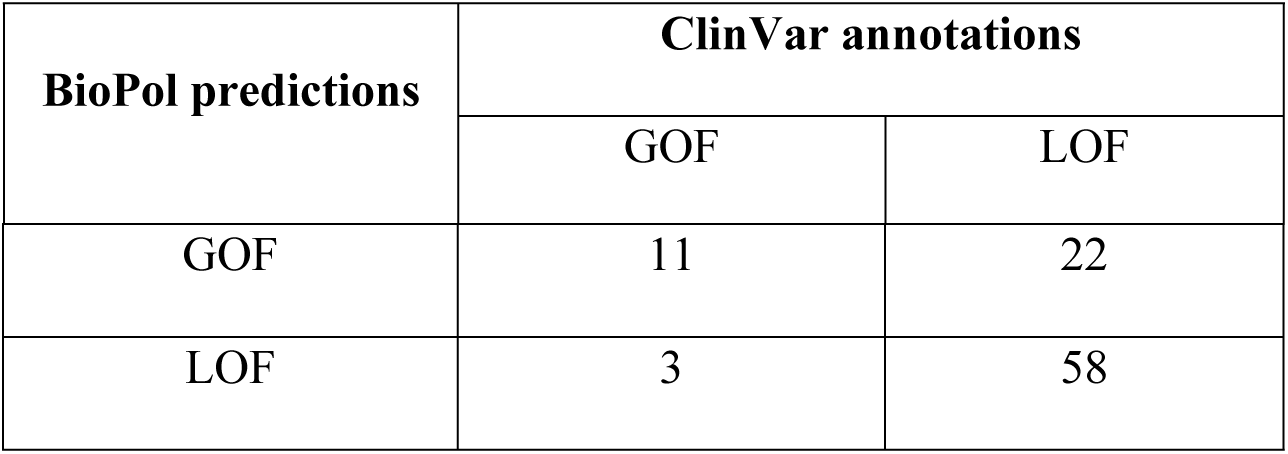
Confusion matrix for BioPol predictions on the validation set.

**Fig 2.**
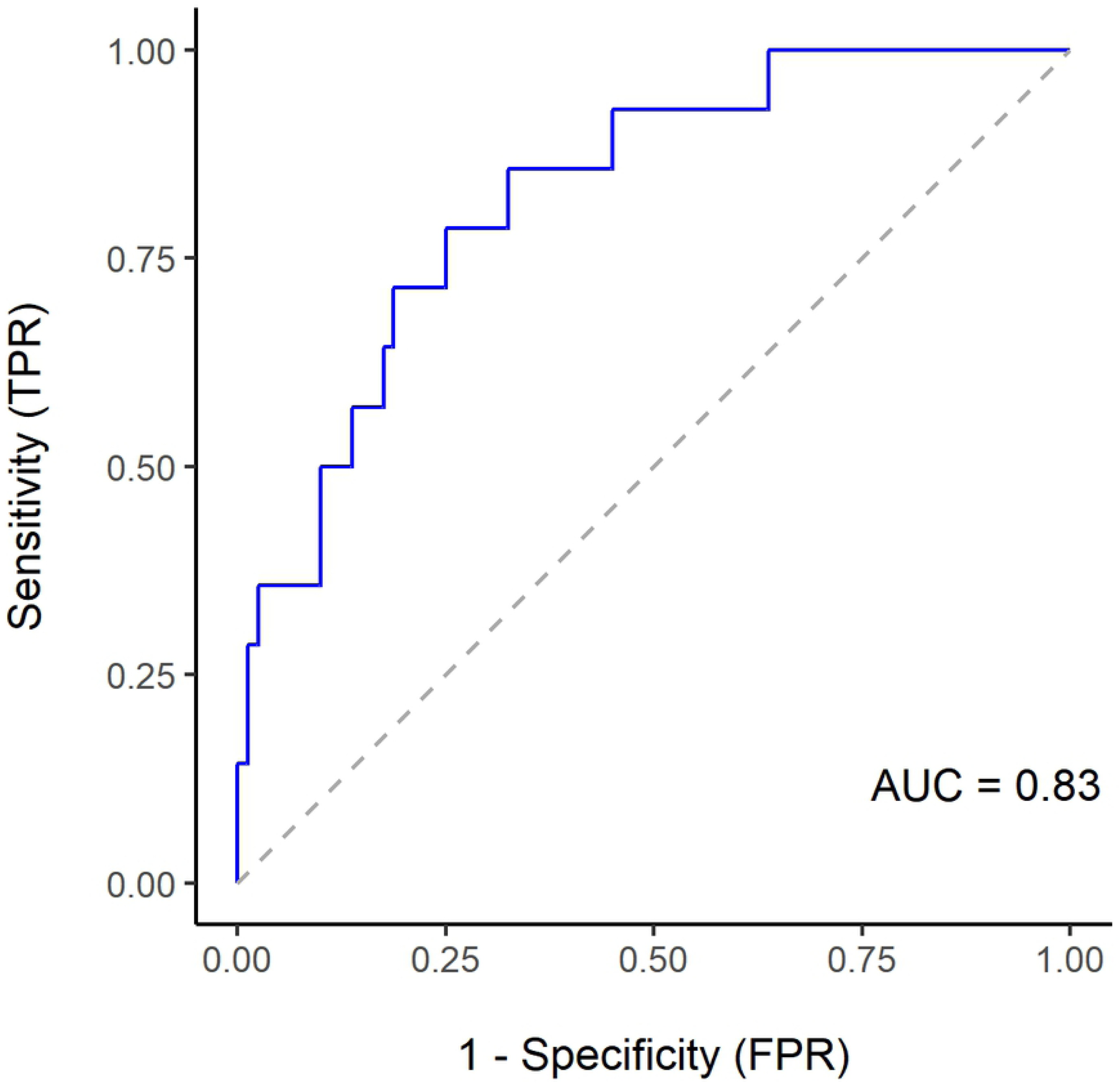
ROC curve illustrating predictive power of BioPol on the validation set.

### BioPol database

We computed BioPol predictions of all possible missense variants that are predicted to be damaging by SIFT, PolyPhen2 and CADD. VEP provides such prediction scores of different tools via dbNSFP [15] plugin module [12]. dbNSFP refers to the database of human non-synonymous SNPs along with their functional predictions [15-16]. Therefore we used it as our primary source for prediction scores. BioPol predictions are stored in a MySQL database and accessible via http://dbe-derm.dbe.unibas.ch/biopol for worldwide users.

## Discussion

Amino acid substitution with large change in physicochemical properties often cause a LOF resulting into protein misfolding. Such large amino acid changes are likely to highly impact the prediction scores of SIFT and PolyPhen2 since they are directly based on sequence and structural features. On the contrary, GOF variants may have a more subtle effect on the protein structure creating a lower impact on SIFT and PolyPhen2 scores. The similar trend of structure-function relationship can also be seen in CADD phred scores as it is one of the key annotations, CADD takes into account while scoring deleteriousness of missense variants.

In conclusion, biological function polarity prediction was found to be possible using the prediction scores of available *in silico* bioinformatics tools SIFT, PolyPhen2 and CADD. Our ensemble classifier BioPol illustrates the same through its high predictive power with AUROC values of >0.8 against two independent sets of damaging variants. However, a weakness of our approach is that the majority of available missense variants have not been investigated with biological experiments. Even though no prediction has perfect accuracy, BioPol predictions can be used as indicators that may help to guide functional experiments on the clinical significance of damaging missense variants.

## Methods

### Missense variant data set

ClinVar seems to be a reliable source for variant data as it archives relationship information between human genetic variants and phenotypic traits with supporting evidences in form of clinical testing reports, research projects and literature texts [11]. Therefore, we extracted the single nucleotide missense variants with GOF and LOF annotations from ClinVar along with their reference human genome assembly GRCh37 locations on 21^st^ September, 2018. We restricted the variants only to pathogenic and likely pathogenic ones based on their clinical significance to maintain high confidence. Also only variants with at least one assertion criteria without any conflicts were selected to increase the confidence level using the review status option of ClinVar [11]. As a result, a total of 298 GOF and 1,583 LOF high confidence variants were obtained.

### Missense variant annotation

The combined set of GOF and LOF missense variants (298 + 1,583 = 1,881) was annotated using the Ensembl web tool VEP [12] (release 96) as it determines the variant effect on genes, transcripts and protein sequences. We restricted the VEP output to one consequence per variant using the flagging option “pick”. The consequences were selected based on a predefined set of criteria including canonical status of transcript, APPRIS isoform annotation, transcript support level, transcript biotype, CCDS status of transcript, consequence rank and transcript length. We filtered out the variants present between two genes i.e., intergenic variants. VEP can provide deleteriousness prediction scores of different *in silico* bioinformatics tools via plugin modules [12]. Hence we extended the annotation output by adding the scores from SIFT, PolyPhen2, CADD, Condel, MutationAssessor, fathmm and MutationTaster.

### Missense variant prediction scores

If a tool was unable to annotate at least 90% of GOF and LOF variants, we excluded it from further analysis. We restricted the annotated variants only to the predicted to be damaging ones by all remaining tools. For each tool, the prediction scores were statistically analyzed using two-tailed unpaired Student’s t-test to check whether they could distinguish between GOF and LOF variants or not. We chose prediction scores of only those tools as biological function polarity predictors, which were significantly different between the two variant classes.

### Missense variant class imbalance

Our variant data set was highly imbalanced as expected, since the number of LOF variants was much higher than that of GOF variants. This directly reflects the natural occurrence as LOF variants are more frequent than GOF ones. In such cases, most standard classification algorithms favor the majority class (LOF) resulting in poor accuracy in the minority class (GOF) prediction. However, many techniques are already available to address class imbalance and can be categorized into data based approaches and algorithm based approaches [17]. Data based approaches handle the class imbalance by providing balanced classes via resampling [18-20]. In our case, resampling would have caused unintentional variant bias. Therefore, we chose gradient boosting algorithm based approach to develop BioPol. Gradient boosting machine sequentially combines weak learners (classifiers) to create strong learners with better accuracy. It starts with the first learner on the training set and calculates the loss. Then it creates an improved learner to minimize this loss function using gradient descent method. This two-step process continues in subsequent iterations until an optimal learner is found or the user defined number of iterations is reached [13].

### BioPol algorithm

We randomly split the final set of missense variants into a training set (70%) and a test set (30%). A seed was set to ensure randomization reproducibility. For a gradient boosting machine model, the main tuning parameters are number of trees or iterations, maximum tree depth and learning rate. We used five separate 10-fold cross validations as resampling scheme to evaluate their effect on model performance and set another seed to control randomness and assure reproducibility. Once the learning rate and maximum tree depth were fixed via resampling, a user defined tuning parameter grid was specified to finalize the number of trees. Equal class weight of 0.5 was added to the model to increase the prediction accuracy of minority GOF class. We chose average AUROC across all cross-validation iterations as the performance measure to select the optimal parameter set and developed BioPol in R using caret [14] package.

### BioPol Performance

Once the classifier was built on the training set, a set of appropriate metrics had to be selected to assess the BioPol predictions on the test set. Overall accuracy can be a misleading metric since the minority GOF class has minimum effect on it. Therefore we selected AUROC as the performance metric [21-22] to evaluate BioPol. Diagnostic plot ROC curve summarizes the discriminative power of BioPol and AUROC is the standard measure for the same. It measures how well GOF and LOF predictions are ranked instead of their absolute values and prediction qualities irrespective of the classification threshold.

### BioPol validation

An independent missense variant set was constructed with GOF and LOF annotations from ClinVar using a different time frame. These variants were submitted to ClinVar between 22^nd^ September 2018 and 25^th^ June 2019. We also ensured that there was no variant overlap between the two data sets. Validation set was similarly filtered for high confidence based on the clinical significance and review status information from ClinVar and annotated using VEP release 96. Only annotated scores of the predictor tools were used to assess how well BioPol can predict the biological function polarity. We used AUROC metric like before to validate the predictive power of BioPol.

### BioPol database

dbNSFP [15-16] version 4.0a aggregates prediction scores of 29 *in silico* bioinformatics tools and hence we used it as the main source for BioPol predictions. We computed predictions of all missense variants that were predicted to be damaging by SIFT, PolyPhen2 and CADD. All of these pre-computed predictions are stored in a MySQL database, hosted at the Department of Biomedical Engineering, University of Basel.

## Acknowledgements

The authors would like to thank Ludovic Amruthalingam and Philippe Gottfrois for their assistance in setting up the website for BioPol database.

## Supporting information

**S1 Table. Set of high confidence GOF and LOF variants with their SIFT, PolyPhen2 and CADD prediction scores**

**S2 Table. BioPol predictions on the validation set of high confidence damaging variants**.

